# Early Neural Correlates of an Auditory Pitch - Visual Size Cross-modal Association

**DOI:** 10.1101/423939

**Authors:** Stephanie C. Boyle, Christoph Kayser, Robin A. A. Ince

## Abstract

Research has shown participants associate high pitch tones with small objects, and low pitch tones with large objects. Yet it remains unclear when these associations emerge in neural signals, and whether or not they are likely the result of predictive coding mechanisms being influenced by multisensory priors. Here we investigated these questions using a modified version of the implicit association task, 128-channel human EEG, and two approaches to single-trial analysis (linear discriminant and mutual information). During two interlaced discrimination tasks (auditory high/low tone and visual small/large circle), one stimulus was presented per trial and the auditory stimulus-response assignment was manipulated. On congruent trials preferred pairings (high tone, small circle) were assigned to the same response key, and on incongruent trials non-preferred pairings were (low tone, small circle). The results showed participants (male and female) responded faster during auditory congruent than incongruent trials. The EEG results showed that acoustic pitch and visual size were represented early in the trial (~100 ms and ~220 ms), over temporal and frontal regions. Neural signals were also modulated by congruency early in the trial for auditory (<100ms) and visual modalities (~200ms). For auditory trials, EEG components were predictive of reaction times, but for visual trials they were not. These EEG results were consistent across analysis methods, demonstrating they are robust to the statistical methodology used. Overall, our data support an early origin of cross-modal associations, and suggest that these may originate during early sensory processing potentially due to predictive coding mechanisms.

## Introduction

Humans exhibit implicit perceptual associations across the different senses. For example, high-pitched tones are often associated with small objects and low-pitched tones with large objects (Gallace and Spence, 2006; Parise and Spence, 2008, 2009, 2012; Evans and Treisman, 2010). These interactions are known as “cross-modal associations”, with preferred pairings defined as “congruent” and non-preferred pairings defined as “incongruent”.

In contrast to multisensory integration, cross-modal associations are situations where information from different sensory modalities interacts with one another, but which does not necessarily result in one percept or a significantly different bimodal neural response (Calvert et al., 2012). However, cross-modal associations have been shown to modulate behavioural performance. For example, participants perform better on congruent compared to incongruent trials (Bien et al., 2012, Parise & Spence, 2008) across a variety of stimulus combinations (Gallace & Spence, 2006). Parise & Spence (2012) even showed that response times were faster for congruent pairings than incongruent pairings, even when only a single stimulus was presented on a trial.

Despite this, where and when associations emerge in neural signals is still unclear (Spence, 2011; Spence and Deroy, 2013; Knoeferle et al., 2017). Bien et al., (2012) examined an auditory pitch-visual size association using an EEG-TMS paradigm, and showed parietal and frontal ERP components were modulated by cross-modal congruency early in the trial (250-300 ms). Kovic, Plunkett, and Westermann, (2010) found early ERP effects at occipital (~140 ms) and parietal (~340 ms) sites which reflected the learned association between semantic (auditory) words and visual objects. Using fMRI, Revill et al., (2014) also showed increased activation in the left superior parietal cortex in response to sound-symbolic pairings of words (e.g. big-slow/small-fast). Finally, Sadaghiani, Maier, and Noppeney, (2009) used fMRI to demonstrate that associations between auditory and visual motion signals emerged in motion areas, whereas higher-level speech-motion associations emerged in fronto-parietal areas.

However, these studies relied on few recording sites (Kovic et al., 2010), sampled brain activity at low temporal resolution (Sadaghiani et al., 2009), or relied on paradigms presenting two stimuli simultaneous. Together this makes it difficult to determine whether stimulus dependent modulations in behavioural responses are due to a genuine cross-modal association or some form of attention dividing or selection (Bien et al., 2012), and as a result it is hard to draw clear conclusions about where and when brain activity reflects cross-modal associations.

In this study we addressed these questions by examining when effects of an auditory pitch – visual size cross-modal association emerged in neural signals. We used the modified version of the implicit association test (IAT) as in Parise & Spence (2012), combined with EEG and two approaches to single-trial analysis. Importantly, using the IAT overcomes the methodological issues mentioned above: it presents only one stimulus per trial (avoiding attentional or multisensory confounds), manipulates congruency by changing the stimulus-response key mapping across blocks (avoiding explicit matching and subjective reports), and presents a single stimulus per trial, which allows us to extract sensory-specific processes from brain activity and relate these to behaviour on a trial-by-trial basis. We hypothesised that reaction times would be faster when congruent pairs of stimuli were assigned to the same response key compared to incongruent pairs. We had two hypotheses regarding the EEG data: a) that brain activity sensitive to the task-relevant sensory feature should be modulated by congruency, and b) that these neural correlates sensitive to the cross-modal congruency should be predictive of participants’ single trial reaction times. We had no prior hypothesis as to when these effects would manifest during a trial or where they would localize; however, if the association arises at an early perceptual level, one would expect their neural correlates to emerge with short latencies after stimulus onset, likely over early sensory areas.

## Methods

### Participants

20 participants (13 females; age range 19-32) took part in the study. One participant’s data (S19) had to be discarded due to noisy EEG channels. The sample size was set to 20, based on sample sizes used in previous EEG studies and general recommendations (Simmons et al., 2011). All participants reported normal or corrected to normal vision and normal hearing. Participants were recruited via the University of Glasgow Subject Pool, and received £6 per hour for their participation. The study was approved by the local ethics committee (application number: 300130001, College of Science and Engineering, University of Glasgow) and conducted in accordance with the Declaration of Helsinki.

### Stimuli

Stimuli were created and presented using MATLAB (MathWorks) and the Psychophysics Toolbox Extensions (Brainard, 1997). Visual stimuli consisted of two light grey circles (‘small’ and ‘large’, 2cm and 5cm, 1.1 ° and 2.8 ° of visual angle respectively) presented for 300 ms atop a darker grey background (Figure 1). Auditory stimuli consisted of two 300 ms pure tones (‘high’ and ‘low’ pitch, 2000Hz and 100Hz respectively). The sound intensity of each tone was matched to 72 Db_A_ SPL (left and right ear) using a sound level meter. Auditory stimuli were presented using Sennheiser headphones and visual stimuli were presented on a Hansol 2100A CRT monitor at a refresh rate of 85 Hz.

**Figure 1.**
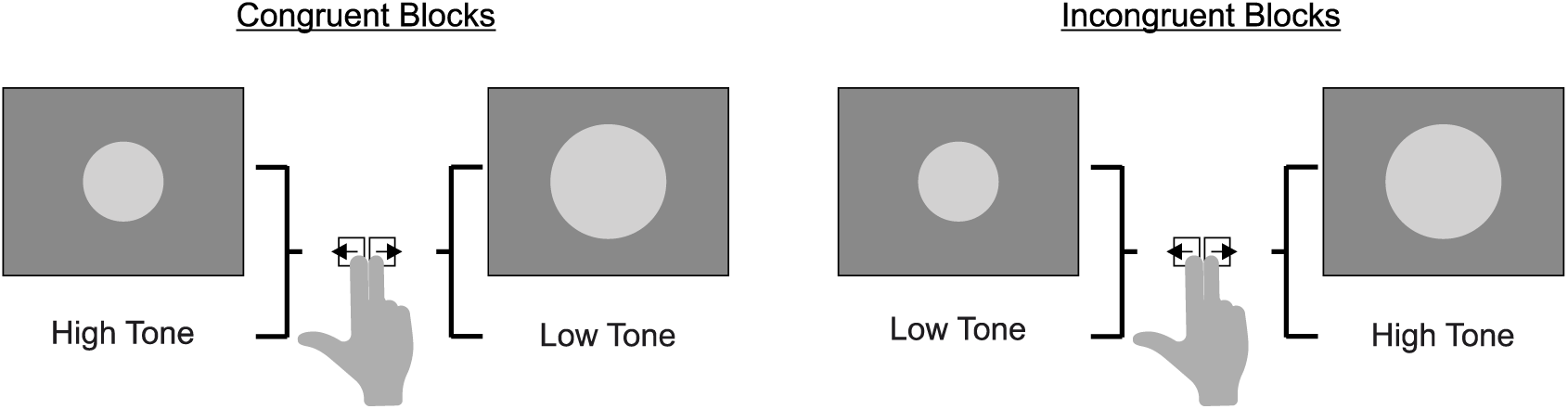
| Task. Subjects were presented with one stimulus (either auditory or visual) on each trial and were asked to indicate which of the two stimuli within that modality was presented (e.g. small vs large circle, high vs low tone) as quickly and accurately as possible. On each block, the assignment of the auditory features to the two response keys was manipulated (*left*, congruent block pairings; *right*, incongruent block pairings).

### Task

The task was a modified version of the IAT (Greenwald et al., 1998), as used in Parise & Spence, (2012). The IAT measures implicit associations via manipulating stimulus-response key mapping, and assumes response times will be faster when associated pairs of stimuli are assigned to the same response key compared to when associated pairs are assigned to different response keys. In this modified version, on each block, one auditory and one visual stimulus are assigned to the left response key, and one auditory and one visual stimulus are assigned to the right (two stimuli per key, see Congruency section for assignment). Participants are then presented with one stimulus per trial, and have to identify which was presented as quickly and accurately as possible using the appropriate response keys. In this particular experiment, cross-modal associations are then measured via reaction time modulations.

### Congruency

Congruency was manipulated by changing the stimulus-response key pairings across blocks (Figure 1). On congruent blocks the small circle and high tone were assigned to the left response key and the large circle and low tone were assigned to the right key. On incongruent blocks the small circle and low tone were assigned to the left response key and the large circle and high tone were assigned to the right response key. Importantly, the auditory assignment changed across blocks, while the visual assignment always remained fixed. In total, participants completed 8 blocks (4 congruent and 4 incongruent presented in a randomised order) for a total of 1280 trials (160 trials per block, 40 trials for each stimulus type).

In this experiment, the stimulus response mapping from one block to the next remained constant for the visual stimulus and only changed for the auditory stimulus for two reasons. First, during informal pilot testing of the stimuli, we found that participants also exhibited a cross-modal association between small visual objects and the left keyboard button, and between large visual objects and the right keyboard button. In order to ensure we did not have confounding effects between associations, we decided to hold the visual stimuli constant and manipulate only the auditory stimuli. Second, each experimental session was approximately three hours long (~2 hour for EEG set up/clean up, and 1.5 hours for the task of 1280 trials). These issues made it difficult to design an implicit association experiment where the visual and auditory stimuli were counterbalanced, and the participants were not asked to spend more than three hours in the lab per session. For these reasons, we chose to only manipulate auditory stimulus congruency.

### Procedure

The experiment was carried out in a dark and electrically shielded room. Each block began with instructions on the mapping between stimuli and response keys (see Congruency). Participants were given as much time as they needed to memorise the instructions for the upcoming block. Each trial started with a fixation cross presented centrally for a randomised period (uniform distribution in 500 to 1000 ms). Then one of the four stimuli (see Stimuli) was selected randomly, and presented for 300 ms (Figure 1). Participants had to respond as quickly as possible using the left and right keyboard keys, as defined by the block instructions (see Congruency). Feedback was provided after each trial (green fixation cross for correct answers, red fixation cross for incorrect answers) for a randomised duration (uniform distribution from 300 ms to 600 ms).

### EEG Recording and Preprocessing

EEG data was recorded using a 128-channel BioSemi system and ActiView recording software (Biosemi, Amsterdam, Netherlands). Signals were digitised at 512 Hz and band-pass filtered online between 0.16 and 100 Hz. Signals originating from ocular muscles were recorded from four additional electrodes placed below and at the outer canthi of each eye.

Data from individual blocks were preprocessed in MATLAB using the FieldTrip toolbox (Oostenveld et al., 2011) and custom scripts. Epochs of 2 seconds (-0.5 to 1.5 seconds relative to stimulus onset) were extracted and filtered between 0.5 and 90 Hz (Butterworth filter) and down-sampled to 200 Hz. Potential signal artefacts were removed using independent component analysis (ICA) as implemented in the FieldTrip toolbox (Oostenveld et al., 2011), and components related to typical eye blink activity or noisy electrode channels were removed. Horizontal, vertical and radial EOG signals were computed using established procedures (Hipp & Siegel, 2013; Keren, Yuval-Greenberg, & Deouell, 2010) and trials where there was a high correlation
between eye movements and components in the EEG data were removed. Remaining trials with amplitudes exceeding ±120 μV were also removed. Successful cleaning was verified by visual inspection of single trials.

## Analysis Methods

### Analysis of Behavioural Data

For each participant, overall performance (proportion correct) and median reaction time (RT) were calculated separately for each modality and congruency. Trials with very fast (<300 ms) or slow (>1200 ms) responses were excluded. Both RT and performance scores were submitted to Wilcoxon Signed rank tests for analysis (see Statistics). All reported RTs are calculated with respect to stimulus onset.

### Analysis of EEG: Linear Discriminant Analysis

We used single-trial, multivariate linear discriminant analysis (Parra et al., 2005; Sajda et al., 2009; Philiastides et al., 2014) to extract discriminant components related to stimulus type, for each modality separately (high versus low pitch; small versus big circle). Prior to analysis, the data was bandpass filtered between 1 and 30 Hz. Next, the classifier (based on regularised linear discriminant analysis, Philiastides et al., 2014), was applied to the EEG activity in sliding time windows of 30 ms, at each 5 ms time point in the window from -300 ms pre-stimulus onset to 1 second post stimulus onset. The discriminant output (Y) was always aligned to the onset of the 30 ms window. Classification performance (Az) was determined using the receiver operator characteristic (ROC) and 10 fold cross-validation along with randomisation testing (see Statistics). Scalp topographies showing the normalised correlation between the discriminant output and the EEG activity were estimated via the forward model (Philiastides et al., 2014).

To assess how these neural correlates of sensory evidence were modulated by congruency, the discriminant output (Y) for congruent trials was compared to that for incongruent trials using a cluster randomisation technique (see Statistics). To examine the relationship between neural and behavioural data, we used linear regression to investigate whether the information contained in the discriminant component (Y) was predictive of behavioural reaction times. As we were interested in whether the quality of the sensory information reflected by the EEG component (i.e. the distance from zero, regardless of sign) was predictive of auditory reaction times, the discriminant output (Y) was flipped for trials that had been assigned to a negative value during classification (e.g. stimulus labels assigned to 0 were multiplied by -1). The discriminant output (Y) for each trial was then regressed against individual participant reaction times at each time point during the trial. Significance levels for both congruency and regression analyses were calculated using a cluster randomisation technique (see Statistics).

### Analysis of EEG: Mutual Information

To assess how robust our effects were, we performed a complementary analysis based on mutual information (MI) (Gross et al., 2013; Ince et al., 2015, 2017; Kayser et al., 2015). MI can be thought of as a likelihood ratio test for dependence between two variables of interest (e.g. between stimulus type and EEG, or between EEG and reaction times). A particular advantage of MI is that it provides a common meaningful effect size scale (bits) across a wide range of statistical tests (Ince, Giordano, et al., 2017).

We calculated MI using a semi-parametric estimator: Gaussian Copula Mutual Information (GCMI) (Ince, Giordano, et al., 2017). This provides a data-efficient and robust lower bound approximation to MI by modelling the dependence between the variables with a Gaussian copula. However, no assumption is made on the marginal distributions of the variables. To allow direct comparison of the results obtained from MI analysis and the linear discriminant analysis, we follow the same pre-processing steps described in the previous section. EEG data were band-pass filtered between 1 and 30 Hz, and then averaged in sliding windows of 30 ms from stimulus onset to 545 ms post stimulus onset in steps of 5 ms (with the data aligned to the onset of the 30 ms window). For each sensor we calculated the single trial central finite difference temporal derivative of the filtered EEG signal. At each time point of each trial, we added this temporal derivate to the voltage to obtain a bivariate response. GCMI allows us to estimate MI using this bivariate response. When a biphasic evoked potential is modulated by an experimental condition, considering voltage alone can often result in a double peak statistical effect, because of the zero crossing where the modulation changes sign (e.g. at the zero crossing of an amplitude modulated bi-phasic waveform). Since MI is an unsigned effect size this results in two positive peaks separated by a time period in which there is no significant effect. Including the EEG voltage and its instantaneous rate of change at each time point addresses this, since at the zero crossing ongoing modulation of the signal can be detected in the gradient. This gives a more balanced picture of the time window over which the EEG signal is modulated by the experimental condition (Ince, Giordano, et al., 2017). GCMI was computed between stimulus type (high/low tone and small/large circle) and the 2D EEG voltage response (EEG data, temporal derivative) for each modality, time point and electrode separately. Statistical significance was determined using randomisation analysis (see Statistics).

To examine when this MI statistic was affected by cross-modal congruency, for each modality we compared MI values (about high/low tone and small/large circle) computed from the congruent trials to MI values obtained from the incongruent trials at each time point. Significant clusters were determined using a cluster randomisation procedure (see Statistics). To examine the relationship between neural activity and behavioural responses, the EEG data in these significant clusters was then averaged over all significant electrodes and a 30 ms epoch over the centre cluster peak. This shorter epoch was chosen as averaging EEG activity over long (>100 ms) windows can cause problems with biphasic evoked potentials due to cancellation of positive and negative periods of the signal. Based on our filtering parameters, we chose a 30ms window around the peak to reduce noise and avoid including periods of signal with different sign.

The resulting single-trial EEG data in these clusters was then regressed against the single-trial reaction times for each participant using multiple linear regression (with each cluster as a predictor). Significance was determined by applying a permutation based t-test across participants on the group level regression weights for each time point, and significant clusters determined using a cluster randomisation technique (see Statistics).

### Statistics

All Z values reported were generated from a two-sided Wilcoxon signed rank test after testing assumptions of normality (which did not hold). Effect sizes were calculated by dividing the Z value by the square root of N (where N = the number of observations rather than participants) (Rosenthal, 1994). P-values were checked for inconsistencies using the R software package “statcheck” (Nuijten et al., 2016).

Statistical significance of classification performance (Az) were determined by randomly shuffling the condition allocated to each trial 2000 times, computing the group averaged Az value (area under ROC curve, see Methods) for each randomisation, and taking the maximal Az value over time for each randomisation. This built a distribution of Az values from which we extracted the 99^th^ percentile. Because of the maximum operation, this provides a Family-Wise Error Rate (FWER) of p = 0.01, corrected for the multiple comparisons over time points (Holmes et al., 1996; Nichols and Holmes, 2001).

Significance for group-level effects of congruency on the discriminant output (Y) were obtained by comparing congruent and incongruent trials across participants at each time point using a permutation based paired t-test across participants (shuffled participant labels, 1000 permutations). Significant clusters were then determined using a cluster randomisation technique which compared the true t-value (resulting from comparing congruent to incongruent Y signals using true participant labels) to the shuffled t-value, based on a cluster threshold of t = 1.8 (p<0.05), maxsum cluster forming, minimum cluster size of 2, and cluster p-value = 0.05. Effect sizes were indicated as the equivalent r value that is bounded between 0 and 1 (Rosenthal & Rubin, 2003).

Significance levels for the LDA regression analysis were generated by randomly shuffling the trial specific reaction times and performing the regression analysis (between the decoding signal and reaction times) 1000 times. Significance was determined by applying a t-test (against zero) across participants on the group level regression weights for each time point, and significant clusters determined using a cluster randomisation technique (with cluster thresholds set as described above).

Finally, statistical significance for the MI analysis was calculated using a randomisation test together with the method of maximum statistics (Holmes et al., 1996). For each time point, sensor, condition, and participant, GCMI was calculated 1000 times with permuted stimulus class labels. The maximum MI value over electrodes and time across permutations was calculated, and the 99^th^ percentile used as the threshold for significance. MI values computed from congruent and incongruent trials were then compared at each electrode and time point using the same cluster randomisation technique (and settings) described above.

## Results

### Behavioural Results

Figure 2 presents the behavioural results. In line with our hypothesis median reaction times were shorter for congruent versus incongruent trials (auditory congruent to incongruent: 959 ms to 993 ms, visual congruent to incongruent: 898 ms to 929 ms, calculated from stimulus offset). As expected, this difference was significant only for auditory stimuli (Wilcoxon sign rank tests: auditory, Z = -2.1328, p = 0.033, effect size = -0.3372; visual, Z = -1.14487, p = 0.127, effect size = -0.229). Performance score did not significantly differ between incongruent and congruent trials (Wilcoxon sign rank tests: auditory, median change congruent to incongruent, 94.3% to 94.1%, Z = -0.402, p = 0.688, effect size = -0.064; visual, median change congruent to incongruent, 97.4 *%* to 97.4%, Z = 0, p = 1, effect size = 0.002).

**Figure 2.**
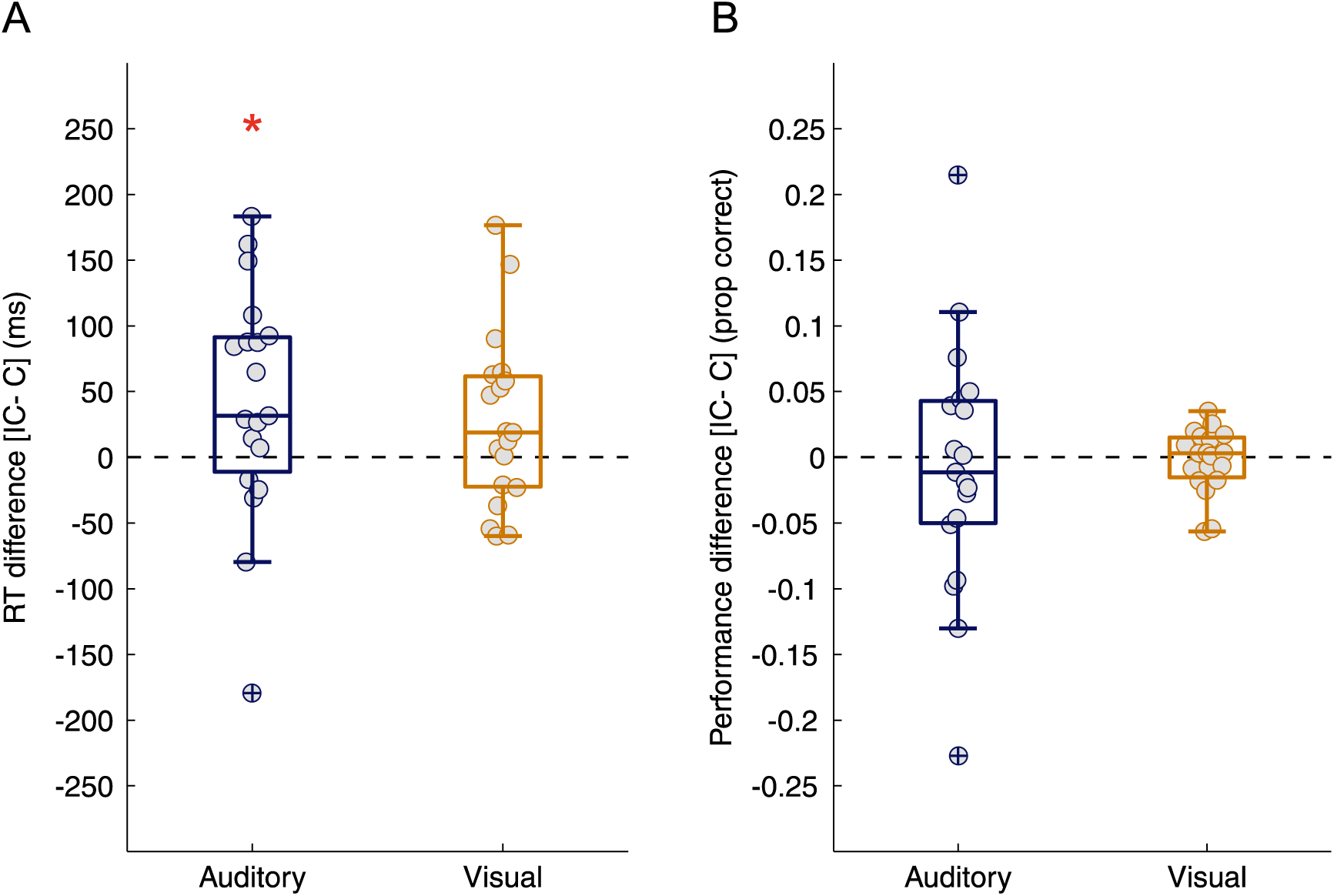
| Behavioural Results. **(A)** Median reaction time difference (Incongruent RT – Congruent RT) across all trials shown for each participant (grey circles). **(B)** Accuracy Difference (Incongruent – Congruent proportion correct). Asterisk (^∗^) represents significant difference (p<0.05, Wilcoxon Signed Rank test).

### EEG Decoding

Figure 3-1 A displays the discriminant performance for auditory and visual trials. For auditory stimuli, significant performance emerged between 25ms after stimulus onset and 535 ms (Figure 3-1 A, blue horizontal dotted line, cluster permutation test, p<0.01, Az value = 0.5387). The corresponding scalp models obtained from the correlation between the discriminant output and the EEG data (averaged over a 30 ms time window centred on the two classification performance peaks) revealed the strongest effects originated over posterior, central and temporal electrodes for both peaks (Figure 3-1 A, topographical inserts, top row). These two topographies were very similar but of opposite sign. For visual stimuli, significant decoding performance emerged between 40ms after stimulus onset and 660 ms (Figure 3-1 A, yellow horizontal dotted line, cluster permutation test, p<0.01 Az value = 0.5432). The corresponding scalp models (Figure 3-1 A, topographical inserts, bottom row) showed strongest correlations between the discriminant output (Y) and the EEG activity over posterior and frontal regions (for the first classification peak and second classification peak respectively). These results indicate that the linear classifier could identify EEG components carrying significant taskrelevant sensory information, and that these components were different for auditory and visual trials.

**Figure 3.1.**
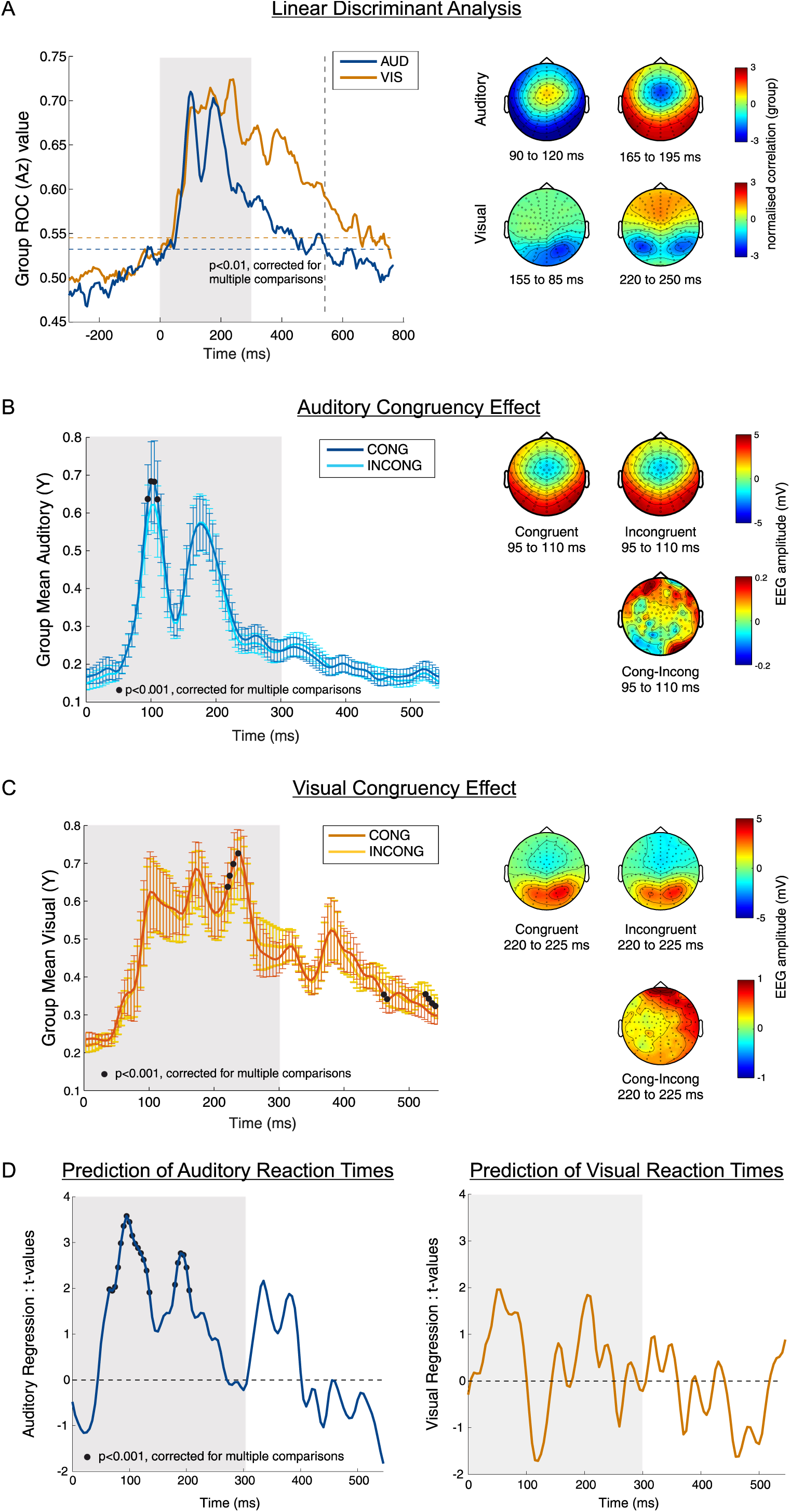
| EEG Linear Discriminant Analysis. **(A)** Group averaged performance of a linear classifier discriminating between stimulus type (auditory: high vs. low tone; visual: small vs. large circle). Auditory classification performance shown in blue, visual classification shown in orange. Horizontal dotted lines represent the threshold for statistical significance (FWER p<0.01, blue for auditory, orange for visual). Vertical dotted line represents the last time point (545 ms) both auditory and visual discrimination are significant (used as a time window cut-off for following analysis). Scalp topographies display the forward models (correlation between discriminant output Y and underlying EEG activity) for a 30ms time window centred on each peak in classification performance (auditory first peak: 105ms, second peak: 180 ms; visual first peak: 170 ms, second peak: 235 ms). **(B)** Group averaged discriminant output (Y) for auditory trials separated by congruency. Black dots represent significant congruency effect (p<0.05, cluster randomisation test). Error bars represent standard eror of the mean. Scalp topographies represent the group-averaged raw EEG activity underlying this significant time window of interest (averaged across time) for congruent and incongruent trials separately. Bottom scalp topography (group averaged, time window averaged, raw EEG data) shows the difference in raw EEG activity (Congruent – Incongruent) **(C)** Group averaged discriminant output (Y) for visual trials, separated by congruency, with significant time points again denoted with black circles. Error bars represent standard eror of the mean. All scalp topographies as in (B), but for the first cluster showing significant differences in the visual modality. Note: we have only included the topographies underlying the first cluster here as this was the only one which occurred during stimulus presentation. To see scalp topographies for the final two significant clusters (cluster 2, 3 occurring after stimulus offset), see Figure 3-2. **(D)** Group-level regression weights (beta values) for the regression of single trial RTs against the discriminant output for auditory (left) and visual (right). Significant time points represented with black circles (p<0.05, cluster permutation test). In all figure panels, grey background represents the time period of stimulus presentation (0 ms to 300 ms).

**Figure 3.2.**
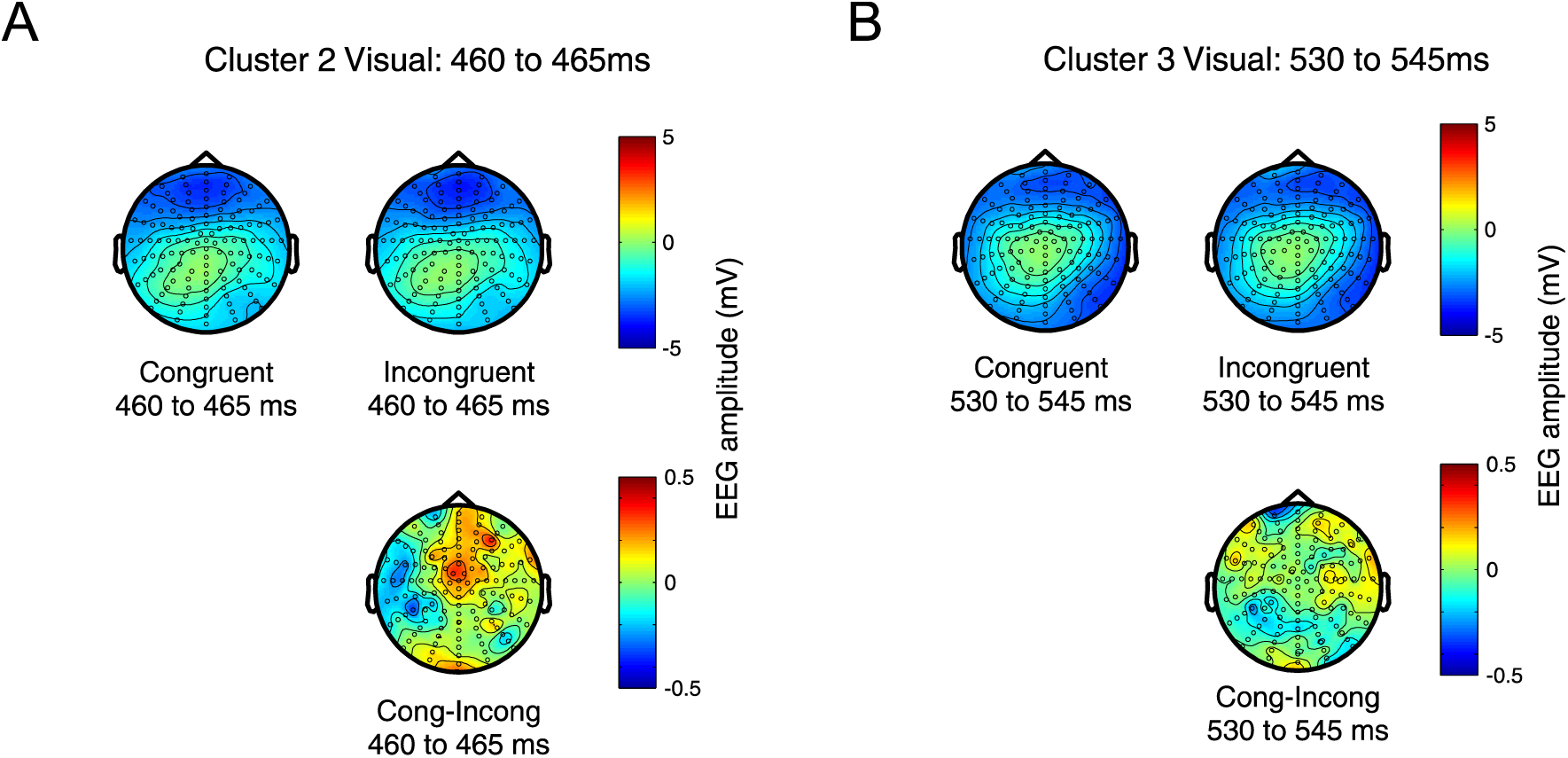
Significant visual LDA clusters (after stimulus offset). **(A)** Scalp topographies represent the group-averaged raw EEG activity underlying the second significant cluster (cluster 2) where congruency differences were found, averaged across time for congruent and incongruent trials separately. Bottom scalp topography (group averaged, time window averaged, raw EEG data) shows the difference in raw EEG activity (Congruent – Incongruent). **(B)** Shows the same as (A) for the third significant visual cluster.

Figure 3-1 B (left) shows the discriminant output (Y) for the auditory modality, separately for congruent and incongruent trials (with the signal Y flipped in order to map all trials to positive values, *see* Methods). For the auditory modality we found one cluster where the discriminant output (Y) was significantly different for congruent compared to incongruent trials (cluster randomisation test, cluster 1: t = 8.576, p<0.001, effect size = 0.4599). This emerged early in the trial, from 95 ms to 110 ms after stimulus onset, and revealed there was more information (i.e. higher value decoding signal) about stimulus type in the congruent signals compared to incongruent signals. The corresponding scalp topographies represent the raw EEG amplitude underlying this time window of interest, shown separately for congruent (Figure 3-1 B, top row, left) and incongruent trials (Figure 3-1 B, top row, right). These revealed similar activation patterns over posterior-temporal regions for auditory congruent and incongruent trials. However, examining the difference (congruent-incongruent) between these raw EEG topographies (Figure 3-1 B bottom row) revealed positive differences over left frontal regions, and right temporal and posterior regions.

Figure 3-1 C (left) displays the discriminant output (Y) for the visual modality, separately for congruent and incongruent trials (again, with the Y signal flipped to map all trials to positive values, *see* Methods). There were three clusters where the discriminant output was modulated by congruency. The first emerged from 220ms to 235ms (cluster randomisation test, cluster 1: t(18) = -7.761, p<0.001, effect size = 0.426). The corresponding scalp topographies displaying the raw EEG activity underlying this time window of interest revealed strong occipital activity for both congruent and incongruent trials (Figure 3-1 C top row, left and right respectively). Examining the difference between congruent and incongruent trials revealed strong positive differences over central frontal and right temporal electrodes. The second cluster emerged after stimulus offset at 460ms to 465ms (cluster randomisation test, cluster 2: t(18) = 4.0906, p<0.001, effect size = 0.441). At this cluster, there was strong negative activity over fronto-central electrodes for both congruent and incongruent trials (Figure 3-2 A, top row, left and right respectively). Examining the difference (congruent – incongruent) between these topographies showed a localised central effect (Figure 3-2 A, bottom row). Finally, the third significant cluster which showed differences based on visual congruency occurred from 530ms to 545ms (cluster randomisation test, cluster 3: t(18) = 9.038, p<0.001, effect size = 0.480). This time, the cluster was associated with strong negative activity over right frontal and temporal electrodes (Figure 3-2 B, top row, left and right respectively). Comparing the scale of the topoplots between auditory and visual conditions (Figure 3-1 B and 3-1 C) shows the early significant auditory condition difference is smaller in amplitude than the later visual one. The similar scale of group mean Az values, the mean and standard error of the discriminant filter Y and the within condition topologies suggests a broadly equivalent signal to noise in terms of stimulus discrimination across the two modalities. The difference in congruency between modalities could be due to higher trial-to-trial variability of the congruence effect, or due to the different congruency mechanisms, either across modalities or early vs later. Finally, examining the EEG signal difference between congruency conditions again revealed-a weaker positive difference over frontal regions between congruent and incongruent trials (Figure 3-2 B, bottom row).

Figure 3-1 D (left) displays the results generated from regressing the auditory decoding signal against auditory trial reaction times. Here we found that the discriminant EEG signals were significantly predictive of reaction times at two clusters during stimulus presentation: from 65 ms to 135 ms, and from 180 ms to 205 ms (cluster randomisation tests: first cluster, t(18) = 40.479, p<0.001, effect size = 0.541; second cluster, t(18) = 14.538, p<0.001, effect size = 0.504). Figure 3-1 D (right) displays result from regressing the visual decoding signal against visual trial reaction times. Here, we found that visual discriminant signals were not predictive of reaction times at any point in the trial (cluster randomisation test, p>0.01).

To summarise the LDA analysis, the results indicated possible differences in the contributions of EEG activity to sensory discrimination and congruency due to the varying time windows and locations of effects. They also demonstrated that congruency effects emerge early during the auditory trials (during stimulus presentation) yet later during visual trials (around stimulus offset and closer to the behavioural response). Finally, the results showed that auditory discriminant EEG signals are predictive of behavioural response times, whilst visual signals are not.

### Mutual Information

Figure 4 A shows the results from the mutual information analysis, for auditory and visual trials separately. For auditory trials, stimulus information (high/low tone) was represented in the EEG signal early in the trial, with significant MI values emerging from 50 ms to 245 ms after stimulus onset (Figure 4 A, blue horizontal dotted line, randomisation test, 99^th^ percentile). This information was highest over left posterior and temporal electrodes (Figure 4 A, topographical inserts, top row). For the visual trials, stimulus information (small/large circle) was represented in the EEG signal early in the trial, with significant MI values emerging from 60ms to 545ms for incongruent trials (Figure 4 A, yellow horizontal dotted line, randomisation test, 99^th^ percentile). This time, the highest information was centred over right posterior electrodes (Figure 4 A, topographical inserts, bottom row). Note that as MI is an unsigned quantity the signs of the values are different, but the spatial patterns obtained from the sensor-wise MI analysis are very similar to the absolute value of the patterns obtained through the LDA forward model (Figure 3-1 A). Figure 4-B (left) displays the results from the congruency comparison for auditory trials, with congruent and incongruent trials shown separately. This revealed stronger MI between the EEG signal and stimulus for congruent compared to incongruent trials, emerging early in the trial. The scalp topographies show that the highest MI for both congruent and incongruent trials occurred over left posterior and right temporal electrodes (Figure 4-B, topographical inserts, top row, MI averaged over a 30 ms time window centred on peak). Comparing MI values between congruent and incongruent auditory trials revealed three significant spatio-temporal clusters: one emerging from 65 ms to 145 ms over frontal electrodes (cluster randomisation test, cluster 1: t(18) = 160.1086, p =0.004, effect size = 0.4176), one emerging 65 ms to 195 ms over left posterior electrodes (cluster randomisation test, cluster 2: t(18) = 180.649, p = 0.004, effect size = 0.4389), and one from 230 ms to 275 ms over right temporal electrodes (cluster randomisation test, cluster 3: t(18) = -131.843, p < 0.001, effect = 0.4751). To avoid artifactual effects in the scalp topographies occurring a result of averaging over these long significant cluster time windows, we reduced the window length of the significant clusters to a 30 ms time window centred on each MI cluster (cluster 1 centre, 105ms; cluster 2 centre, 130ms; cluster 3 centre, 250ms). Figure 4-B (topographical inserts, bottom row) displays the early differences found over frontal (bottom left), temporal (bottom middle), and posterior (bottom right) electrodes for each shorter spatio-temporal cluster (Figure 4 B, topographical inserts, bottom row).

**Figure 4.**
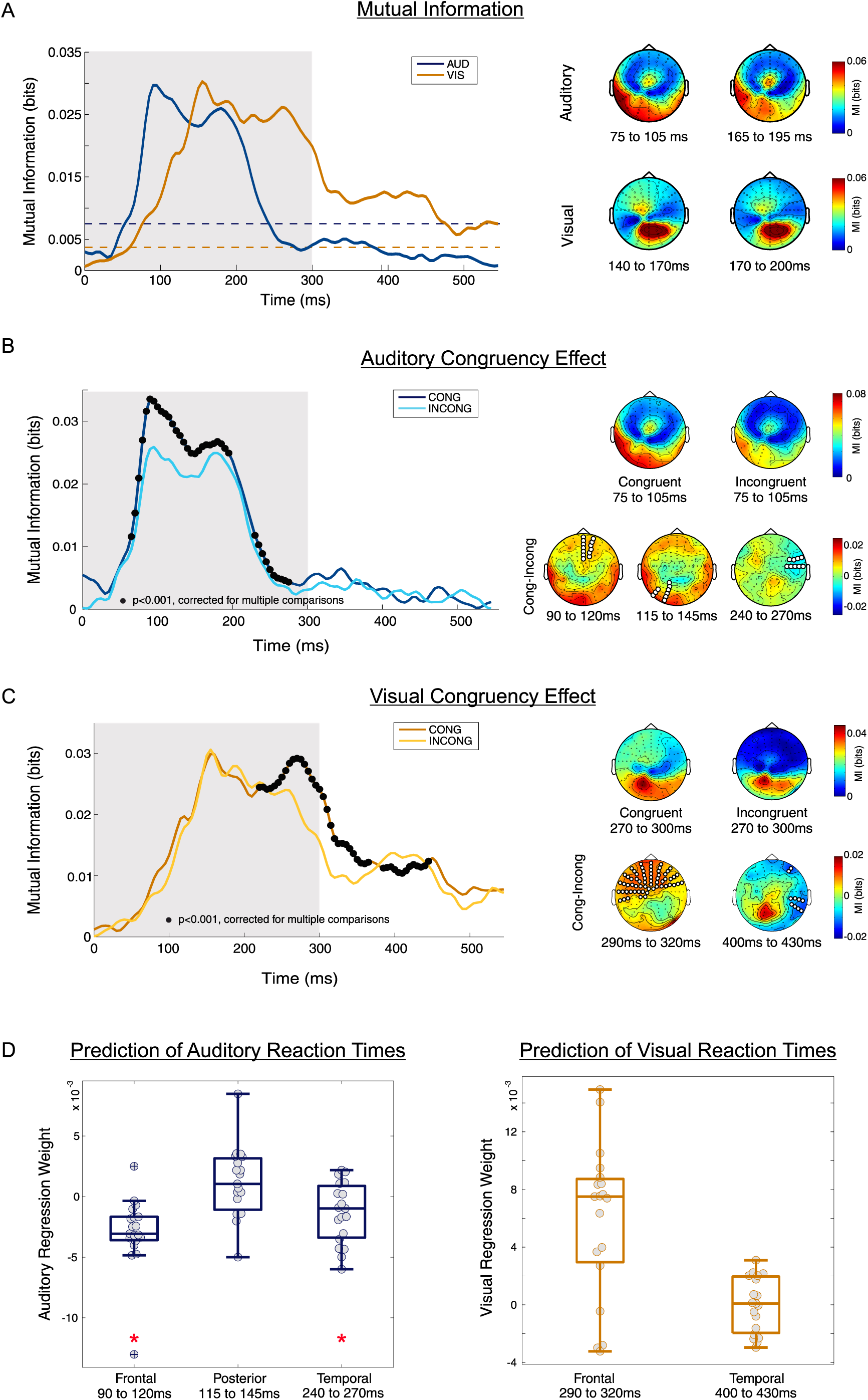
| EEG Mutual Information Analysis. **(A)** Group level mutual information (MI) between EEG and stimulus type (auditory: high vs. low tone, visual: small vs. large circle) averaged over electrodes. Auditory MI shown in blue, visual MI shown in orange. Dotted lines show time windows where the MI for each condition was significant (blue dotted line = auditory significance, orange dotted line = visual significance). Scalp topographies (right) show the MI at peak time points for auditory (top row) and visual (bottom row) separately. **(B)** Congruency difference between congruent and incongruent auditory MI (averaged over electrodes). Significant time points where there was a difference between congruent and incongruent MI represented with black circles (p<0.05, cluster randomisation test). Scalp topography (B, top row) shows MI underlying congruent (left) and incongruent (right) auditory trials underlying peak MI (averaged over 30ms around centre of peak MI difference). Scalp topography (B, bottom row) shows the MI difference (Congruent – Incongruent) for the three clusters where there was a significant congruency effect (again, topographies are activity averaged over 30ms around centre of each significant cluster). **(C)** Same as in (B), but for visual trials. **(D)** Regression weights (beta values) generated from regressing EEG activity against reaction time for the three clusters of interest for auditory (left) and visual (right) trials. Grey circles represent individual subject beta weights, and asterisks represent clusters where EEG activity significantly predicted reaction time. Again, in all panels grey background represents the time of stimulus presentation (0 ms to 300 ms).

Figure 4 C (left) displays the results from the congruency comparison for visual trials, with congruent and incongruent trials shown separately. Again, this revealed stronger MI between the EEG signal and stimulus for congruent compared to incongruent visual trials. The scalp topographies indicated that MI between the EEG signal and stimulus type was strongest over left posterior electrodes and frontal electrodes (Figure 4 C, topographical inserts, top row, MI averaged over a 30ms time window centred on peak). In contrast to the auditory modality, comparing congruent and incongruent visual MI revealed significant differences at two spatio-temporal clusters later in the trial: one from 220 to 365ms (cluster randomisation test, cluster 1 : t(18) = 250.67, p <0.001, effect = 0.4848) over frontal and right temporal electrodes, and one from 385 to 445ms (cluster randomisation test, cluster 2: t(18) = - 99.309, p <0.001, effect = 0.4309) over temporal electrodes. Again, the time window for displaying each spatio-temporal cluster was reduced to a 30 ms time window centred on each cluster which showed significant differences based on congruency (cluster 1 centre, 290ms; cluster 2 centre, 415ms). Figure 4-C (topographical inserts, bottom row) displays the differences found over frontal (bottom left) and temporal (bottom right) electrodes for each shorter spatio-temporal cluster.

Finally, Figure 4-D displays the results of the regression analysis between the EEG activity underlying the significant clusters and single-trial reaction times for auditory (left) and visual (right) separately. As a reminder, to calculate the regression weights we averaged the EEG activity over all significant electrodes (shown in Figure 4-B and 4-C, white circles) and over the shorter, 30ms windows centred over the cluster peaks (defined above) and regressed the activity in each cluster against behavioural reaction times. For auditory trials, this regression analysis demonstrated that the EEG in both the early frontal and later temporal cluster was predictive of reaction times (cluster 1 : 90 ms to 120 ms, t(18) = -4.3073, p = 0.00042, effect size = -0.988; cluster 3: 235 ms to 265 ms, t(18) = -2.2979, p = 0.0338, effect size = -0.5272 for frontal and temporal respectively). However, auditory activity in the posterior cluster was not significantly predictive of reaction times (cluster 2: 115 ms to 145 ms, t(18), = 1.349, p = 0.194, d = 0.309). For the visual trials, the EEG activity in both the frontal and temporal clusters was not significantly predictive of visual trial reaction times.

To summarise, the MI analysis results indicated that the encoding of acoustic information was affected by congruency early in the trial, and activity over frontal and temporal electrodes was significantly predictive of reaction times at these early latencies. In contrast, the encoding of visual information was affected by congruency only later in the trial over frontal and temporal electrodes, and this activity was not significantly predictive of reaction times.

### Consistency of Results

Both EEG analyses revealed that the EEG signal contained information about stimulus type during stimulus presentation, with the highest information emerging over posterior and temporal regions for auditory trials, and over posterior regions for visual trials (see Figure 3-1 A and Figure 4 A). In both analyses, effects of stimulus congruency emerged early in the trial for auditory (Figure 3-1 B and Figure 4 B), and the strongest differences appeared over posterior, temporal and frontal regions. In both analyses, effects of stimulus congruency for visual trials emerged later in the trial with the strongest differences over frontal and temporal regions (Figure 3-1 C and Figure 4 C). Finally, both analyses demonstrated that neural activity was predictive of behavioural reaction time during stimulus presentation for auditory trials, and that neural activity underlying visual trials was not predictive of behavioural reaction time (see Figure 3-1 D, Figure 4 D). Overall, these similar findings across methods are encouraging, and demonstrate that our results are robust. While both methods detect representation of the two stimuli in both modalities with effects well above the threshold for statistical significance, we note that mutual information seems more sensitive to differences in the strength of the effect between congruence conditions, with larger proportional differences in effect size and larger significant clusters. This suggests the particular properties of the mutual information effect size scale might be well suited to this sort of between condition comparison that are common in neuroimaging (Kayser et al., 2015; Park et al., 2015; Keitel et al., 2017).

## Discussion

In this experiment we examined the neural mechanisms underlying an auditory pitch-visual size cross-modal association. Overall, participants responded faster on auditory congruent than incongruent trials. The EEG data showed that stimulus information could be extracted from EEG activity during stimulus presentation for both auditory and visual trials, and that effects related to stimulus type were strongest over posterior and temporal regions respectively. Auditory neural correlates were modulated by congruency early in the trial (<100 ms), and these differences emerged from frontal, posterior and temporal regions. Visual neural correlates were modulated by congruency later in the trial (>250ms) over frontal and temporal areas. Finally, auditory activity underlying a congruency modulated representation of the auditory stimulus in frontal and temporal regions was also significantly predictive of single trial auditory reaction times, indicating that these activations reflect a neural correlate of the underlying perceptual association. In contrast, activity underlying a congruency modulated visual stimulus representation was not significantly predictive of visual reaction times. Importantly, these results were consistent across two separate analysis methods, showing our results are robust.

## Effect of Cross-modal Congruency on Behaviour

The behavioural results showed that participants had faster reaction times for auditory congruent stimulus-response assignments than incongruent ones. This provides further evidence supporting the existence of an acoustic pitch – visual size association, which has been reported in various experimental paradigms before (Bien, ten Oever, Goebel, & Sack, 2012; Evans & Treisman, 2010; Gallace & Spence, 2006; Marks, Ben-Artzi, & Lakatos, 2003; Parise & Spence, 2008; Parise & Spence, 2009; Parise & Spence, 2012). More importantly, our results demonstrate that a cross-modal association between pitch and size can occur even when only a single stimulus is presented on a trial. This finding replicates the work using the IAT carried out by Parise & Spence (2012). Furthermore, as the IAT presents only one stimulus per a trial, it rules out the possibility that our effects are due to general multisensory benefits (i.e. due to spatial and temporal congruency), or to attentional differences caused by dividing attention between two stimuli.

## Early Neural Correlates of Pitch-Size Association

Our results revealed that auditory neural signals were modulated by the congruency of sensory information early during the trial (<100 ms after stimulus onset), while visual trials were modulated later (>220ms). The early onset of these results suggests that these cross-modal auditory associations arise at an early stage, suggesting they are possibly perceptual in origin rather than exclusively decision related. Supporting this interpretation, a recent study by Mostert, Kok, & de Lange, (2015) used MEG, two tasks (one sensory, one sensory and decision making), and a dual decoding approach to demonstrate that sensory related processes emerged in occipital areas from 130 ms, whereas decisional related processes emerged later, around 250 ms. These timings are broadly consistent with our work, with both auditory and visual modulations occurring earlier than 220ms after stimulus onset.

The timing of our results are also broadly in line with previous neuroimaging studies. For example Kovic et al., (2010) found neural signals were modulated around 140 ms to 180 ms by congruency (sound-symbolic association), while Bien et al., (2011) found effects emerging at 250 ms. The short latency onset of these effects occurring in response to the presentation of two associated arbitrary stimulus properties led both sets of authors to conclude that these associations arise in the early stages of the multisensory integration process, in line with past work that has shown multisensory interactions occurring in neural signals at very short latencies after stimulus onset (Giard & Peronnet, 1999; Molholm et al., 2002; Sperdin et al., 2009). However, in our study the early modulations we observe cannot be due to multisensory integration, as on each trial only a single stimulus was presented. We suggest that the congruency effects we observed may be due to some form of top-down perceptual feedback influencing signals in the different modalities at an early stage during the perceptual process.

Alternatively, it could relate to some existing underlying mapping of the perceptual priors of a pitch-size association, which automatically influences early sensory processing. The acoustic pitch – visual size association considered here is strongly reflected in the statistics of the natural world, due to the physics of acoustic resonance (where larger objects resonate at lower frequencies than smaller ones). Given that such cross-modal associations reflect a naturally occurring link between stimuli (Parise et al., 2014), this could lead to a strong Bayesian prior on this relationship (Kersten et al., 2004; Knill and Pouget, 2004; de-Wit et al., 2010; Huang and Rao, 2011). Therefore, one interpretation is that the early sensory effects we observe could be related to influences of such priors in the framework of predictive coding. For example, if caused by top-down signals to early sensory areas, this feedback might embed the environmental prior. If caused by existing underlying mapping in early sensory areas, this suggests that long term environmental priors maybe implemented directly in early sensory areas. Important to note, this particular association may be more intrinsic, innate, and based on real world acoustic properties than other cross-modal associations are (Asano et al., 2015). Consequently, the effects of this particular association as measured by an implicit task may arise earlier in neural signals, than would effects of a semantic association measured via an indirect task.

Finally, we observed that early (<100 ms) neural correlates of auditory congruency were predictive of behavioural reaction times, while visual correlates emerged later (>200ms) and were not. Given that we also find an effect of congruency on behaviour for auditory trials, but not for visual trials, this suggests that the auditory modulations are more behaviourally relevant than the visual. The early auditory modulations could therefore represent the updating of audio-visual congruency, whereas the visual modulations could represent a stable decisional related correlate based on visual congruency. Alternatively, the modulations in auditory signals could represent a very early decision stage, which specifically relates to the mapping between stimulus and response key and arises from the early encoding of congruency during stimulus presentation. In the current paradigm, it is difficult to conclusively prove whether the early auditory modulations arise relate to perceptual process or a decision-making. However, the early onset of the effect (<100 ms after stimulus onset), leads us to speculate that these modulations are potentially perceptual in origin.

Finally, it is important to note that the divergent results between auditory and visual stimuli is almost certainly due to the experimental design whereby the auditory congruency is manipulated while visual congruency is held constant. The results presented here do not mean that cross-modal associations follow different rules in the auditory and visual modalities. Rather, they show how congruency is affected in auditory signals and it can allow us to hypothesise that if the design were reversed (i.e. visual congruency is manipulated and auditory held constant), we would find early modulations in visual signals.

## Spatial Distribution of Early Cross-Modal Effects

Both the LDA and MI analysis revealed that congruency differences emerged over frontal, temporal and posterior areas for auditory trials, and frontal and temporal electrodes for visual trials. These findings are consistent with other studies investigating how neural activity is modulated during cross-modal associations: Bien et al., (2012) found modulations in ERPs over frontal regions, and using fMRI Sadaghiani, Maier, & Noppeney, (2009) demonstrated that higher-level speech-motion cross-modal associations emerged in fronto-parietal areas.

Interestingly, modulations in parietal activity are the most common finding in previous studies examining the neural underpinnings of cross-modal associations. Bien et al., (2012) found that parietal ERPs (200 ms - 300 ms) were modulated by congruency, and that TMS applied over parietal cortex reduced the amplitude difference between congruent and incongruent trials, and the behavioural congruency effect. Similarly, Kovic, Plunkett, & Westermann, (2010) found early ERP modulations over parietal sites, and Sadaghiani, Maier, & Noppeney, (2009) found an interaction in fronto-parietal areas in response to audio-visual motion stimuli. In our experiment, we found that parietal activity was not predictive of behaviour or modulated by congruency, and so our findings are at odds with these previous results. Yet the findings of these previous studies are confounded with the issue of multisensory integration and attention (as described in the introduction), and so parietal activity observed in these previous studies may be reflecting multisensory processing or attention, rather than reflecting a pure cross-modal association. As a result, the effects of parietal TMS on cross-modal effects in the Bien study might simply disrupt multisensory integration processes rather than specific cross-modal association processes, and the parietal activation seen in the Kovic et al., (2010) and Sadaghiani, Maier and Noppeney (2009) study may reflect audio-visual processing or divided attention. In contrast, in our experiment we presented only a single stimulus on each trial, thus ruling out effects of multisensory integration or divided attention. We propose that this suggests the parietal component represents effects due to multisensory integration or attention, and that the effects we describe here may provide a more accurate picture behaviourally relevant effects of cross-modal associations on early sensory processing in the brain.

## Acknowledgements

SCB funded by BBSRC DTP Studentship. BBSRC http://www.bbsrc.ac.uk/ (grant number BB/L027534/1) to CK. European Research Council https://erc.europa.eu/ (grant number 646657) to CK.

